# Alcohol causes lasting differential transcription in *Drosophila* mushroom body neurons

**DOI:** 10.1101/752477

**Authors:** Emily Petruccelli, Nicolas Ledru, Karla R. Kaun

**Affiliations:** Department of Neuroscience, Brown University, Providence, RI, USA, 02912; Department of Biological Sciences, Southern Illinois University Edwardsville, Edwardsville, Illinois, USA, 62026

**Keywords:** alcohol, memory, RNA-seq, splicing, *Drosophila*

## Abstract

Repeated alcohol experiences can produce long-lasting memories for sensory cues associated with intoxication. These memories can ultimately trigger relapse in individuals recovering from alcohol use disorder (AUD). The molecular mechanisms by which alcohol changes memories to become long-lasting and inflexible remain unclear. New methods to analyze gene expression within precise neuronal cell-types can provide further insight towards AUD prevention and treatment. Here, we employed genetic tools in *Drosophila melanogaster* to investigate the lasting consequences of ethanol on transcription in memory-encoding neurons. *Drosophila* rely on mushroom body (MB) neurons to make associative memories, including memories of ethanol-associated sensory cues. Differential expression analyses found that distinct transcripts, but not genes, in the MB were associated with experiencing ethanol alone compared to forming a memory of an odor cue associated with ethanol. These findings reveal the dynamic and highly context-specific regulation of splicing associated with encoding behavioral experiences. Our data thus demonstrate that alcohol can have lasting effects on transcription and RNA processing during memory formation, and identify new transcript targets for future AUD and addiction investigation.

## Introduction

Alcohol use disorder (AUD) impacts millions of individuals and constitutes one of the most serious public health problems in the world today (Kendler *et al*. 2016; Grant *et al*. 2017; Cheng *et al*. 2018). This chronic relapsing brain disorder can persist for extremely long periods of time regardless of alcohol abstinence. Relapse can be triggered by exposure to cues that predict alcohol availability, as they evoke memories of the drug’s effects to result in cravings (Jasinska *et al*. 2014; Courtney *et al*. 2016; Groefsema *et al*. 2016; Valyear *et al*. 2017; Clemens and Holmes 2018; Logge *et al*. 2019). The complexity of alcohol’s molecular actions and our limited understanding of how these actions change distinct neurons in the brain’s reward memory circuitry has prevented our understanding of the mechanisms underlying these maladaptive memories.

Unlike the majority of other abused drugs, ethanol does not act on a single molecular target, but instead affects a variety of molecules (Nestler 2013; Trudell *et al*. 2014). Many of the molecules implicated in ethanol-induced behaviors have broad roles in regulating diverse processes such as cell signaling, transcription, and neuronal plasticity (Ron and Barak 2016; Erickson *et al*. 2019). These roles are more often than not dependent on both cell-type and developmental stage. The complexity of these processes, and diversity of experimental approaches in research studies, often obscures ethanol’s context-dependent molecular effects. However, recent advances in cell-type specific isolation and sequencing technology can reveal the precise consequences of alcohol exposure on gene expression.

The genetic tools available in the fruit fly *Drosophila melanogaster* provide the ability to define where, when, and how alcohol may be acting in the nervous system, including within memory circuitry. *Drosophila* demonstrate ethanol-induced hyperactivity, tolerance (Wolf *et al*. 2002), and consummatory preference (Devineni and Heberlein 2009). They also remember and prefer the experience of intoxication (Kaun *et al*. 2011; Nunez *et al*. 2018), and this process requires mushroom body (MB) neuron activity. The MB integrates both sensory odor information from olfactory projection neurons and valence (aversive/rewarding) information from dopamine neurons. Downstream MB output neurons then drive avoidance of, or approach toward, an odor cue (Aso *et al*. 2014).

Groundbreaking studies have demonstrated that long-term memory formation in many species requires both *de novo* transcription and protein synthesis (see reviews (Bailey *et al*. 1996; Alberini and Kandel 2014; Sweatt 2016). Although long-term memory formation requires transcriptional changes (Alberini and Kandel 2014; Uchida and Shumyatsky 2018), it remains unclear whether repeated ethanol experiences influence these processes. Similarly, whether the presentation of ethanol alone or ethanol paired with an odor induces the same transcriptional events is unknown. We hypothesized that ethanol exposures, alone or paired with an odor, would produce lasting transcriptomic changes within memory-associated MB neurons, thus altering memory formation.

Recent RNA-sequencing (RNA-seq) analysis in *Drosophila* has systematically identified the transcriptomic profiles of various cell-types within MB circuitry (Shih *et al*. 2019), and examined gene expression changes in the context of associative memory formation (Crocker *et al*. 2016; Widmer *et al*. 2018). We used nuclear, MB-specific RNA-seq to better understand transcriptional changes after the formation of alcohol cue associative memories. We found that alcohol exposures caused lasting expression changes at the transcript, but not gene, level, and that altered transcript expression was distinct between alcohol alone and alcohol cue memory groups. This suggests that RNA processing provides a distinct molecular landscape to help encode specific behavioral experiences.

## Materials and Methods

### Fly Husbandry

Flies were raised on cornmeal agar food at 24**°**C, 70% humidity, and with a 14:10 Light:Dark cycle. Male 4-5 day-old flies were isolated under CO_2_, given one day to recover, and then conditioned with specified paradigms detailed below. The fly lines used in this study are a pan-MB specific driver *MB010B* split Gal4 line (Bloomington Stock #68293)(Aso *et al*. 2014) and the 5x*UAS*-*unc84*-2x*GFP* ‘INTACT’ line (Henry *et al*. 2012). Experimental flies were heterozygous for transgenes.

### Odor cue-induced ethanol memory

Odor cue-induced ethanol memory was performed as published previously (Kaun *et al*. 2011; Nunez *et al*. 2018; Petruccelli *et al*. 2018). A minor difference was the use of smaller training/testing vials with 20-30 flies/vial. Humidified ethanol vapor (90:60 EtOH:air), resulting in a moderate 13.8 ± 3 mM (0.01 g/dL) internal body ethanol concentration was used. Ethanol, odors (1:36 odorant:mineral oil with ethyl acetate or iso-amyl alcohol), or both were delivered to flies in perforated vials for 10 min, resulting in a moderate internal ethanol concentration (Kaun *et al*. 2011). Following a 50 min rest, second and third sessions were then performed. Vapor treatments were delivered to flies on 1% agar and supplemented with yeast pellets overnight before being sacrificed the next day by liquid nitrogen freezing. All behavioral experiments were performed in reciprocal to control for innate odor preferences.

For each biological replicate, 2,000 male fly heads were collected via liquid nitrogen freezing and sieve separation 24 hours after conditioning. Flies were conditioned with one of four exposure paradigms (Figure 1A). Biological replicates were performed for Air (*n* = 3), EtOH (*n* = 3), Odors (*n* = 4), and Trained (*n* = 4) conditions. Odors and Trained conditions were performed with reciprocal odor groups (*n* = 2) (Ethyl Acetate→Isoamyl Alcohol and Isoamyl Alcohol→Ethyl Acetate).

**Figure 1.**
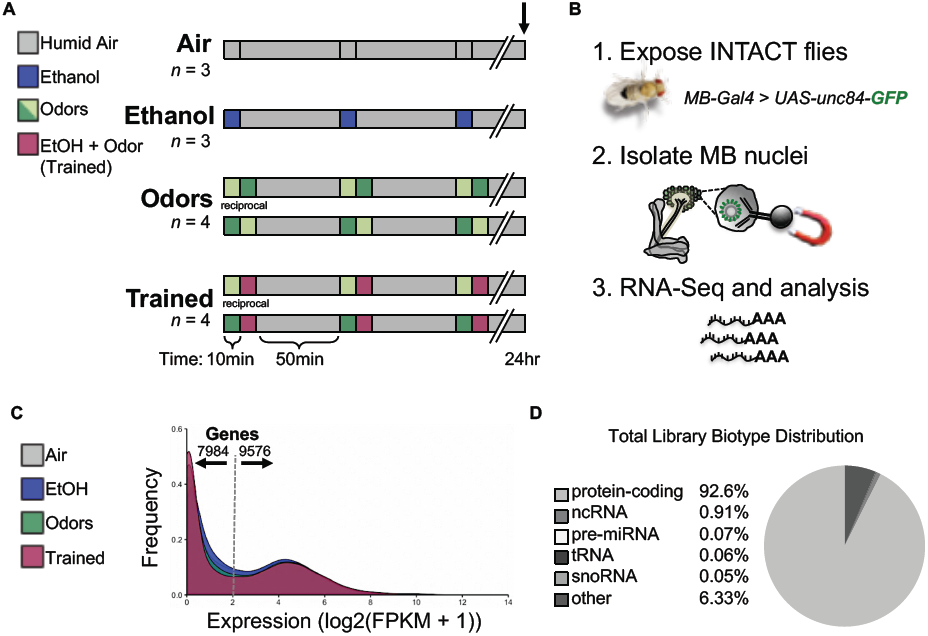
Mushroom Body INTACT paradigm and initial analysis. **A)** Paradigm depicting three spaced 10 minute exposures of ‘Air’, ‘Ethanol’, ‘Odors’, or ‘Trained’ (odor + ethanol) and sacrificed 24 hours later. Each biological replicate (n) was ∼2000 males (Air: *n* = 3, Ethanol: *n* = 3, Odors: *n* = 4 (2 reciprocals), Trained: n = 4 (2 reciprocals). **B)** RNA-seq was performed on mushroom body (MB) nuclei of flies expressing the INTACT transgene (*MB10B-Gal4* > *UAS-unc-84-GFP*). Each fly brain contains ∼4000 MB nuclei. **C)** Histogram of the mean gene expression in fragments per kilobase (FPKM) across treatment libraries. **D)** Percent distribution of biotypes according to current annotated Flybase ‘feature’ types across treatment libraries.

### Isolation of mushroom body nuclei (INTACT procedure)

Isolation of Nuclei Tagged in specific Cell Types (INTACT) was adapted from the method described in (Pankova and Borst 2016) and pioneered in flies by (Henry *et al*. 2012). The INTACT method of extracting nuclear RNA for sequencing provides several advantages compared to a general RNA-seq approach. First, the ability to use flash-frozen tissue, in contrast to FACS or dissected tissue samples, allows for an accurate examination of the current transcriptional profile of genetically targeted cells. Second, nuclear RNA from neurons contributes to the integrity of the active transcriptome snapshot by minimizing contamination from mRNA stored in the cytoplasm, along dendrites, and within axons, while allowing for the detection of experience-dependent differential expression (Lacar *et al*. 2016). Finally, given evidence that mRNA handling in subcellular compartments has been implicated in the formation and storage of memory (Bramham and Wells 2007; Richter 2010; Shigeoka *et al*. 2016; Nakahata and Yasuda 2018; Biever *et al*. 2019), this approach provides a robust profile of the stable post-exposure transcriptome unencumbered by the diversity of whole cell RNA.

Frozen heads were homogenized in a Kontes glass homogenizer (Sigma, D9938-1SET) with 10 mL of dounce buffer (10 mM ß-glycerophosphate, 2 mM MgCl_2_, 0.5% Igepal buffer) and homogenized for ∼2 min with the loose A pestle. Homogenate was passed through a 190 µm nylon net filter (Small Parts, CMN-0185-C). The filter was subsequently rinsed with 2 mL of the same buffer before discarding. Homogenate was subsequently further homogenized gently ∼6-7 times with the tight B pestle, passed through a 20 µm filter (Small Parts, F020N-12-C), brought up to 50 mL by adding sucrose buffer (10 mM ß-glycerophosphate, 2 mM MgCl_2_, 25 mM KCl, 250 mM sucrose). Next, 300 µL of Dynabead A magnetic beads (Thermo, 10001D) were pre-incubated with rabbit α-GFP antibody (Thermo, G10362), and incubated with the homogenate for 30 min at 4°C with gentle agitation. Beads were then captured with a magnet for 15 min and washed 5 times with 600 µL of sucrose buffer for 5 min at 4°C with gentle agitation. Finally, RNA was extracted from bead-bound nuclei using TRIzol (Ambion, Life Technologies), resuspended in RNA-ase free water and DNA-ase treated according to manufacturer’s instructions (Ambion DNA-Free Kit).

### RNA-seq

RNA libraries were polyA enriched to identify mRNA transcripts ready for nuclear export and sequenced using a Hi-Seq 4000 (Illumina) machine at a depth of ∼30 million single-end 1×50bp reads by GENEWIZ (South Plainfield, NJ). Read quality was assessed using FastQC-0.11.5 (Babraham Bioinformatics). Adapters were removed and trimmed using Trimmomatic-0.36 (Bolger *et al*. 2014). The ‘new tuxedo suite’ was used to further process reads (Pertea *et al*. 2016). Reads were aligned with HISAT2-2.0.5 (Pertea *et al*. 2016) to the Ensembl BDGP6_transcriptome reference (Dm6), modified to include the *unc-84-2xGFP* sequence present in our flies. Next samtools-1.3.1 (Li *et al*. 2009) and stringtie-1.3.3 (Pertea *et al*. 2016) were used to sort, merge, and quantify transcripts. Lastly, assembly and analysis of Ballgown objects (Pertea *et al*. 2016) and data visualization with ggplot2 was performed in RStudio-1.0.136 (Wickham 2009). No samples were discarded and default settings were used for the pipeline unless otherwise noted - command line code as follows:

~~~
*#Trimmomatic:*
TruSeq3-SE.fa:2:30:10:8:true LEADING:3 TRAILING:3 SLIDINGWINDOW:4:15 MINLEN:36
*#HISAT2:*
hisat2 -q -x <*reference index*> -U <*input fastq file*> -S <*output sam file*> -p 50
*#Samtools:*
samtools view -Su alns.sam | samtools sort -o alns.sorted -@ 4
*#StringTie:*
stringtie <*input*.*sorted*.*bam*> -b <*path to Ballgown tables*> -o <*output*.*gtf*> -G <*reference*.*gtf*>
~~~

### Statistical Analysis

Libraries were statistically analyzed in R to determine differential expression between treatment conditions. Raw p-values are shown, as well as corrected p-values (false discovery rate < 0.05). The freely available Ballgown package (Pertea *et al*. 2016) afforded general linear modeling statistics for differential expression analysis and data visualization tools. Libraries were normalized with Ballgown package default – sum of each library’s log expression measurements below the 75th percentile (Frazee *et al*. 2015). For intersectional analyses, hypergeometric statistics were performed on pairwise comparisons and considered different from chance at p-value < 0.05.

## Results

### RNA-sequencing library characterization

We first generated flies that express the nuclear-tagged GFP ‘INTACT’ transgene (*UAS-unc84-GFP*) (Henry *et al*. 2012) in all MB neurons (*MB010B-Gal4*). Expression of INTACT was MB specific and did not to disrupt alcohol cue memory formation (Supplemental Figure 1). MB-INTACT flies were subjected to one of four treatment paradigms – Air, EtOH, Odors, or Trained and flash frozen the following day (Figure 1A). MB nuclei were isolated and RNA sequenced to a depth of around 30 million reads per sample (Supplemental Figure 2). Nearly all libraries showed >80% uniquely mapped reads and ∼15 million FPKM (Fragments Per Kilobase of transcript per Million mapped reads) per sample (Supplemental Figure 2).

A distribution of mean expression across the 17560 annotated genes revealed a similar frequency distribution of gene expression across treatment conditions (Figure 1C). Across the entire dataset, 7984 genes (45%) showed little to no expression (<2 log2(FPKM+1), whereas 9576 genes (55%) had robustly detected levels of gene expression (≥2 log2(FPKM+1). This suggested that adult MB nuclei were actively transcribing roughly 55% of annotated genes in the genome at the time of isolation.

The BDGP6 Dm6 transcriptome is comprised of 17560 genes and 34741 transcript isoforms with 10010 genes uniquely represented by a single transcript isoform. According to the current Flybase (flybase.org) annotation information, the Dm6 contains roughly 85% protein-coding isoforms, 2% noncoding RNAs, 1.5% microRNAs, 0.9% tRNAs, 0.8% snoRNAs, and 9.7% other/non-annotated biotype sequences. When pooling all expression counts across treatments, our data was comprised of 92% protein-coding genes/isoforms, 0.9% noncoding RNAs, 0.06% microRNAs, 0.06% tRNAs, 0.05% snoRNAs, and 6.3% other/non-annotated sequences (Figure 1D). Together the characterization of our RNA-sequencing libraries suggests that adult MB neurons are actively transcribing just over half of the genes in the genome and that there is an abundance of protein-coding genes detected to allow for interesting comparisons between treatment conditions.

### Expression of *a priori* and novel gene lists across treatments

Since our RNA-seq approach was restricted to MB nuclei, we expected to see prominent expression of genes previously associated with MB function. Using the ‘Vocabularies’ search function on FlyBase (flybase.org), we acquired a list of genes associated with the ‘adult mushroom body’ (FBbt:00003684) including *Octopamine receptor in mushroom bodies* (*Oamb*), *Protein kinase cAMP-dependent regulatory subunit type 2* (*Pka-R2*), *short Neuropeptide F precursor* (*sNPF*), *Vesicular Acetylcholine Transporter* (*VAChT*), *portabella* (*prt*), *Dopamine Ecdysone Receptor* (*DopEcR*), etc. As expected, the majority of these MB-associated genes showed robust mean expression (>2 log2(FPKM+1)) across our treatment libraries (Figure 2A) thus supporting the MB specificity of the libraries.

**Figure 2.**
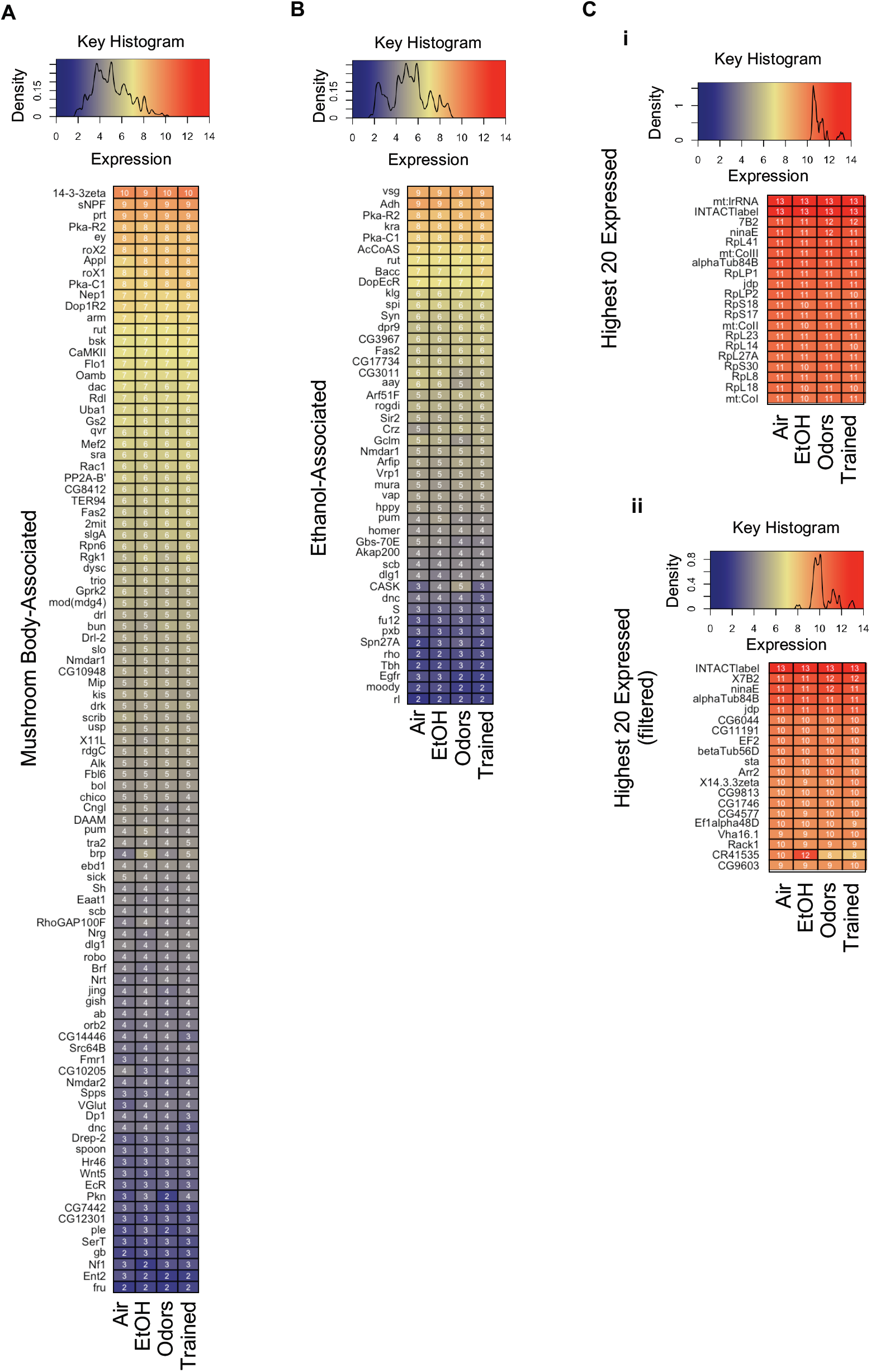
Highest expressed genes across behavioral conditions. Density histogram and keys with blue, yellow, and red color representing low, medium, and high expression for **(A)** MB-associated, **(B)** ethanol-associated, and **(C)** highest expressed genes. All gene expression represented as condition means (log2(FPKM+1)) and ordered by expression. **C)** The highest 20 genes expressed **(i)** without filtering, and **(ii)** after a rough filtering of RNA-associated genes.

Since the treatment conditions differed in the presence or absence of ethanol, we investigated how many ethanol-associated genes were expressed in MB nuclei. The acquired FlyBase gene list for ‘behavioral response to ethanol’ genes (FBbt:0048149) included *Sirtuin 1* (*Sirt1*), *Epidermal growth factor receptor* (*Egfr*), *homer* (*homer*), *Alcohol dehydrogenase* (*Adh*), *happyhour* (*hppy*), etc (Figure 2B). Forty-six of the fifty-five ethanol-associated genes showed robust expression (>2 log2(FPKM+1)) in the MB, but few genes showed noticeable differences between treatment conditions. This was not surprising, however, since most ethanol-associated genes were discovered using high-dose, acute ethanol treatment throughout the whole fly or heads, while this data shows gene expression 24 hours after repeated mild ethanol exposures specific to MB nuclei.

We next investigated the highest 20 transcribed genes across treatments (Figure 2C). As expected, we found the ‘*INTACTlabel’* (manually added to the Dm6 transcriptome) to be highly expressed across the libraries, demonstrating the robust nature of the *GAL4/UAS* binary expression system and further validating our MB isolation technique.

Aside from the INTACT label, most of the highly transcribed genes were RNA-associated genes (Figure 2Ci), which we posit to be a mix of transcripts from rough endoplasmic reticulum that was pulled down during nuclear isolation, as well as newly transcribed or nuclear-localized RNAs. Previous researchers using INTACT may not have observed these findings because ribosomal proteins were masked during transcriptome alignment (Henry *et al*. 2012). Theoretically, nuclear RNA-seq should not require extensive ribosomal depletion and may allow for interesting rRNA expression analyses.

We then curated the highest 20 mean expressed genes after excluding RNA-associated genes (Figure 2Cii). These included genes encoding rhodopsin *NinaE* and rhodopsin inactivation protein (*Arr2*), translation elongation factors *EF1alpha48D* and *EF2*, a PKC inhibitor (*14-3-4Z*), a vacuolar ATPase (*VHA16-1*), and cytoskeleton associated proteins *alphaTub84B, betaTub66D,Rack1* and others. Together, these descriptive analyses indicate that our libraries represent comparable MB datasets in which lasting experience-dependent molecular effects could be explored.

### Experience-dependent differential gene expression

To uncover experience-dependent effects on transcription, we first examined genes that showed the greatest variance in expression (log transformed counts per million means) across treatments. This descriptive statistic reveals those genes with the greatest variability due to experience (variance is the square of the standard deviation from the mean). Among the 100 most variable genes, many ribosomal proteins and previously uncharacterized (unnamed CG#) genes were identified. Also included were *Esterase-6* (*Est-6*), *Cytochrome P450-4g1* (*Cyp4g1*), *la costa* (*lcs*), *Mucin related 18B* (*Mur18B*), *alpha and beta Trypsin (βTry, βTry*), *yippee interacting protein 7* (*yip7*), *fab1 kinase* (*fab1*), *Troponin C (TpnC73F and TpnC47D*), *Odorant binding protein 51a* (*Obp51a*), *obstructor-B* (*obst-B*), *inaF-B* (*inaF-B*), *Autophagy-related 10* (*Atg-10*) *short spindle 2* (*ssp2*), *neurotransmitter transporter-like* (*Ntl*), *Multiple inositol polyphosphate phosphatase 2* (*Mipp2*), and a number of *Jonah* peptidase transcripts (Figure 3Ai). Excluding RNA-associated genes further revealed *Maltase Ai* (*Mal-A1*), *Sarcolamban B* (*SclB*), *Cuticular protein 49Ab* (*Cpr49Ab*) and *Coiled-coil domain containing 56* (*Ccdc56*) (Figure 3Aii).

**Figure 3.**
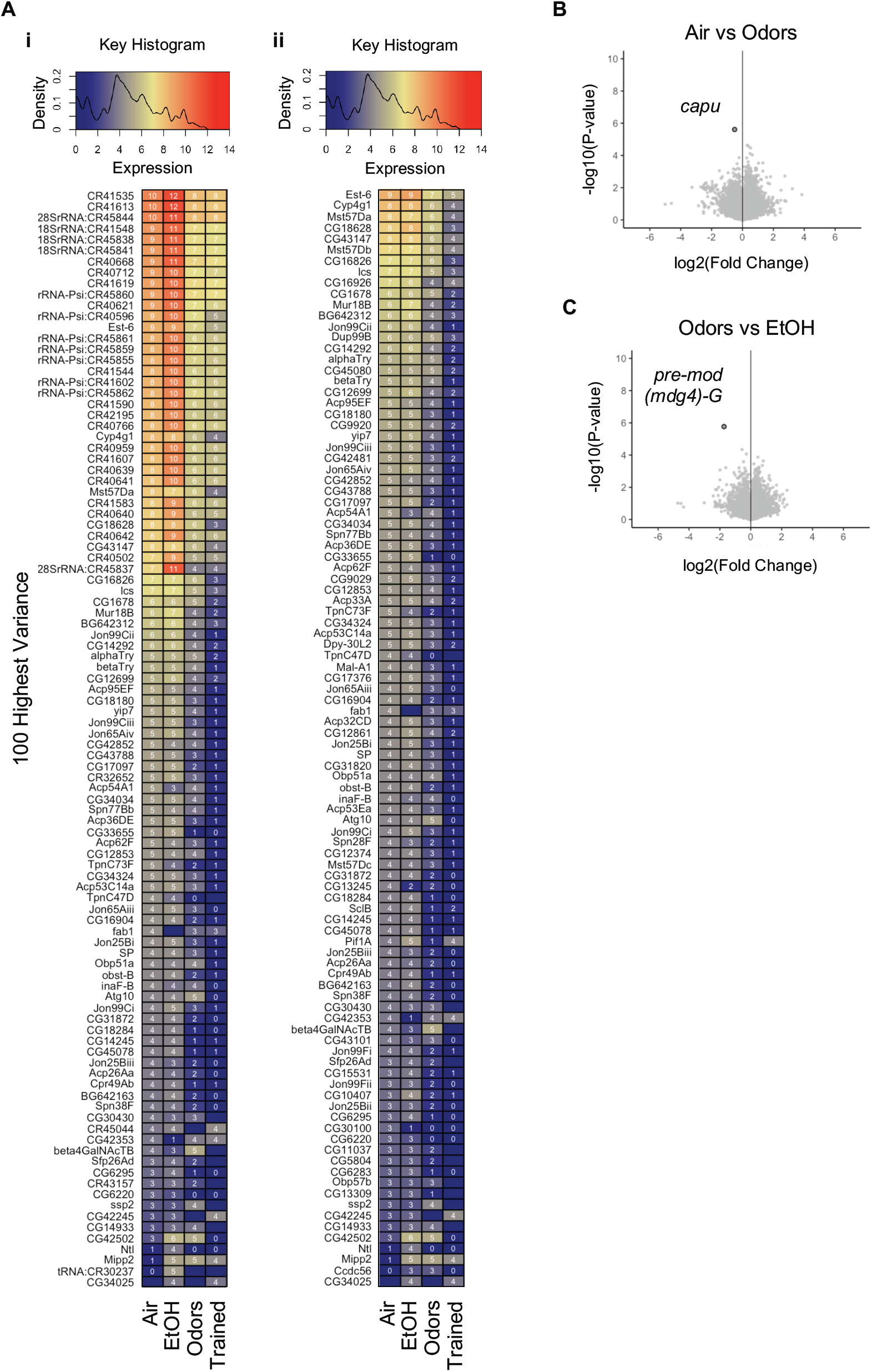
Top most variable and statistically significant genes between treatments. **A)** Density histogram and keys with blue, yellow, and red color representing low, medium, and high expression of the 100 most variable genes – the squared deviation from the mean – **(i)** without filtering, **(ii)** after a rough filtering of RNA-associated genes. **B-C)** Volcano plot showing gene fold change (log2(fc) compared to the inverse of significance (-log10(p-value) (dark outline, FDR < 0.05). Both **(B)** Air vs Odors and **(C)** Odors vs EtOH had one statistically significant gene that changed expression.

*Sex Peptide* (*SP*), *Male-specific RNA 57Da and b* (*Msdt57Da, Msdt57Dd*), *BG642312, Serpins 77Bb* and *38F, Ductus ejaculatorius peptide 99B* (*Dup99B*), *Dpy-30-like 2* (*Dpy-30L2*), *PFTAIRE-interacting factor 1A* (*Pif1A*), *Seminal fluid protein 26Ad* (*Sfp26Ad*), and a number of *Accessory gland protein* (*Acp*) transcripts were also found, but may be a consequence of using male flies as these proteins are known to be produced by the male accessory glands. However, since heads were isolated prior to nuclear extraction, and the highly specific MB010B split-Gal4 line is not known to be expressed in the male accessory gland (Aso *et al*. 2014), our findings may be explained by a few contaminating whole fly bodies in some samples, or be unappreciated bona fide transcripts in MB neurons.

To determine differential expression between treatment conditions, we used ‘Ballgown’ (Pertea *et al*. 2016), a Bioconductor package. No genes or transcripts were removed from analyses and the adjusted p-value (a.k.a. q-value) was set to the standard false discovery rate (FDR) < 0.05. All pairwise condition contrasts were examined at both the gene and transcript level (Supplemental Figure 2). Only two genes were differentially expressed 24 hours after the various treatments, the actin filament nucleating protein *cappuccino (capu)* (Air vs Odors) (Figure 3B) and the nuclear DNA binding protein pre-*mod(mdg4)-G* (Odors vs EtOH) (Figure 3C).

### Intersectional analysis of differential isoform expression

In contrast to gene-level analysis, there were more statistically significant differences found at the transcript level despite being subjected to more repeated measure correction. This suggests that there must be altered transcriptional start/stop site or RNA processing events that altered isoform diversity. In light of this finding, we focused on the differential expression of transcript isoforms between treatments.

Using Air treatment as a control, we compared the intersection of the top 200 most significantly differentially expressed (lowest p-values) isoforms from Air vs Odors (Figure 4Ai), Air vs EtOH (Figure 4Aii) and Air vs Trained (Figure 4Aiii). There was surprisingly little overlap in specific isoforms, including between ‘Air vs EtOH’ and ‘Air vs Trained’ (Figure 4B). This indicated that experiencing odors, ethanol intoxication, or making ethanol-odor memories produced distinct, molecularly separable changes in the lasting transcriptional state of MB nuclei. Using hypergeometric statistics, the number of overlapping isoforms was found to be statistically greater than would be expected by chance alone (Supplemental Figure 3). The extent of intersection was visualized with an upgraded Venn Diagram plot generated by an R package called ‘UpSetR’ (Figure 4B). Overlapping top p-value isoforms between treatment conditions are listed in Figure 4C.

**Figure 4.**
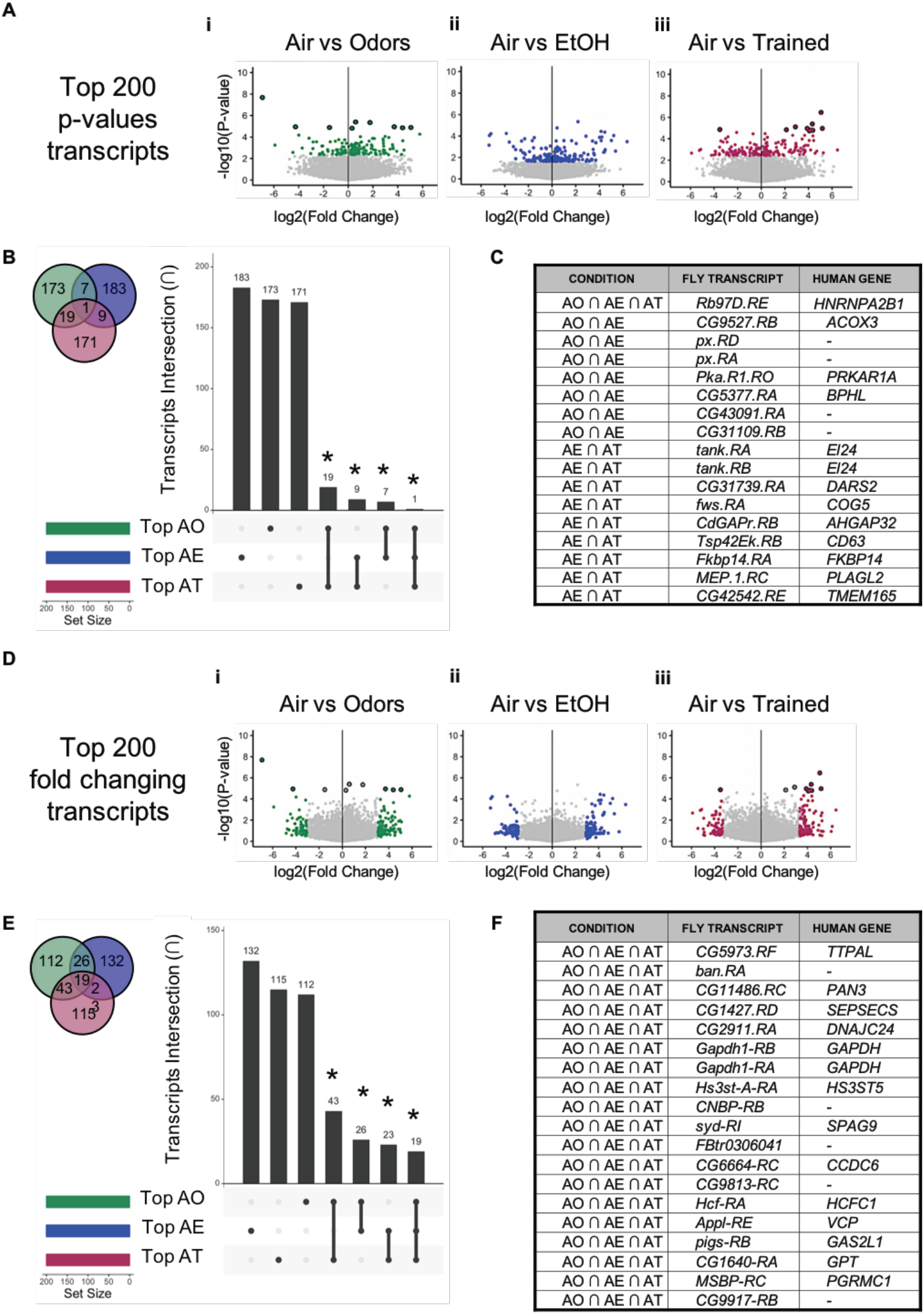
Intersectional analysis of top differentially expressed transcripts across treatments, as compared to Air Controls. **A)** Volcano plots of **(i)** Air vs Odors, **(ii)** Air vs EtOH and **(iii)** Air vs Trained comparisons (dark outline, FDR < 0.05). Colors depict the top 200 p-value transcripts. **B)** An upgraded Venn Diagram plot generated by an R package called ‘UpSetR’ demonstrating intersection in top 200 p-value transcripts (abbreviated by first letter in treatment). **C)** A few of the transcripts from intersectional analysis in **(B)** are listed in a table along with corresponding high DIOPT scoring human genes. **D)** Volcano plots of **(i)** Air vs Odors, **(ii)** Air vs EtOH and **(iii)** Air vs Trained comparisons. Colors depict the top 200 fold changing transcripts. **E)** An UpSetR plot demonstrating intersection in top 200 fold-change differentially expressed transcripts. **F)** A few of the transcripts from intersectional analysis in **(D)** are listed in a table along with corresponding high DIOPT scoring human genes.

Using the same logic just described, we compared the 200 isoforms with the largest fold changes in expression (Figure 4D). This approach highlights whether specific isoforms have greater dynamic range in expression in response to experiencing odors, ethanol, or ethanol-odor conditioning. Interestingly, there were more overlapping isoforms between pairwise comparisons than observed in the top p-value transcripts intersection analysis (Figure 4E). This finding suggests that particular isoforms in the MB neurons are more plastic than others to being changed in response to Odors, EtOH, and Trained treatments. Hypergeometric statistics again showed that the number of overlapping isoforms was statistically greater than would be expected by chance alone (Supplemental Figure 3). A subset of the overlapping top fold-changing isoforms between treatment conditions are listed in Figure 4F. The intersectional analyses between all pairwise library comparisons can be found in Supplemental Figure 4.

### Enrichment of genes associated with differential isoforms

To determine which types of isoforms are enriched in the top 200 p-value comparisons, we performed DAVID Gene Ontology (GO) analysis on the associated gene IDs for each treatment - 167 Air, 175 Odor, 175, Ethanol, and 188 Trained genes. Using default DAVID settings, the top three GO annotations were identified (Supplemental Figure 5). Compared to Air controls, repeated Ethanol exposures produced statistically significant GO terms – ‘Alternative splicing’ (p-value = 0.0002), ‘Phosphoprotein’ (p-value = 0.041), and ‘Coiled Coil’ (p-value = 0.047). Interestingly, the response to Trained treatment also had ‘Alternative splicing’ (p-value = 0.062) as a high scoring enrichment category. It is important to note that the ‘Alternative splicing’ GO term does not refer to spliceosome genes, but rather to genes that are ‘known to be alternatively spliced’. The enrichment analysis results further support the idea that there are experience-dependent changes in transcriptional start/stop site regulation or RNA processing, but not overall gene level changes. Perhaps future ontology databases will include isoform-specific categorization.

### Candidate transcripts underlying ethanol cue memories

This study was motivated by our interest in identifying whether lasting gene expression changes are associated with formation of ethanol cue memories. As our goal was to investigate how ethanol affected gene expression in memory circuits during memory formation, we were most intrigued by the comparisons between Odor vs Ethanol, Ethanol vs Trained, and Odor vs Trained flies.

We found seven transcripts (six genes) that were differentially expressed in response to Ethanol treatment as compared to Odors only controls (Figure 5A). One transcript was downregulated – *CG17982*^*-RA*^ – and six were upregulated – translational regulator *Dodeca-satellite-binding protein 1* (*Dp1*^*-RH*^), transcription factor *elbow B* (*elB*^*-RD*^), transcription factor *modifier of mdge 4* (*premod(mdg4)-G*^*-RA*^), hypoxia-induced protein *CG11825*^*-RA*^, and genes of unknown function *CG17982*^*-RB*^, and *CG17982*^*-RA*^.

**Figure 5.**
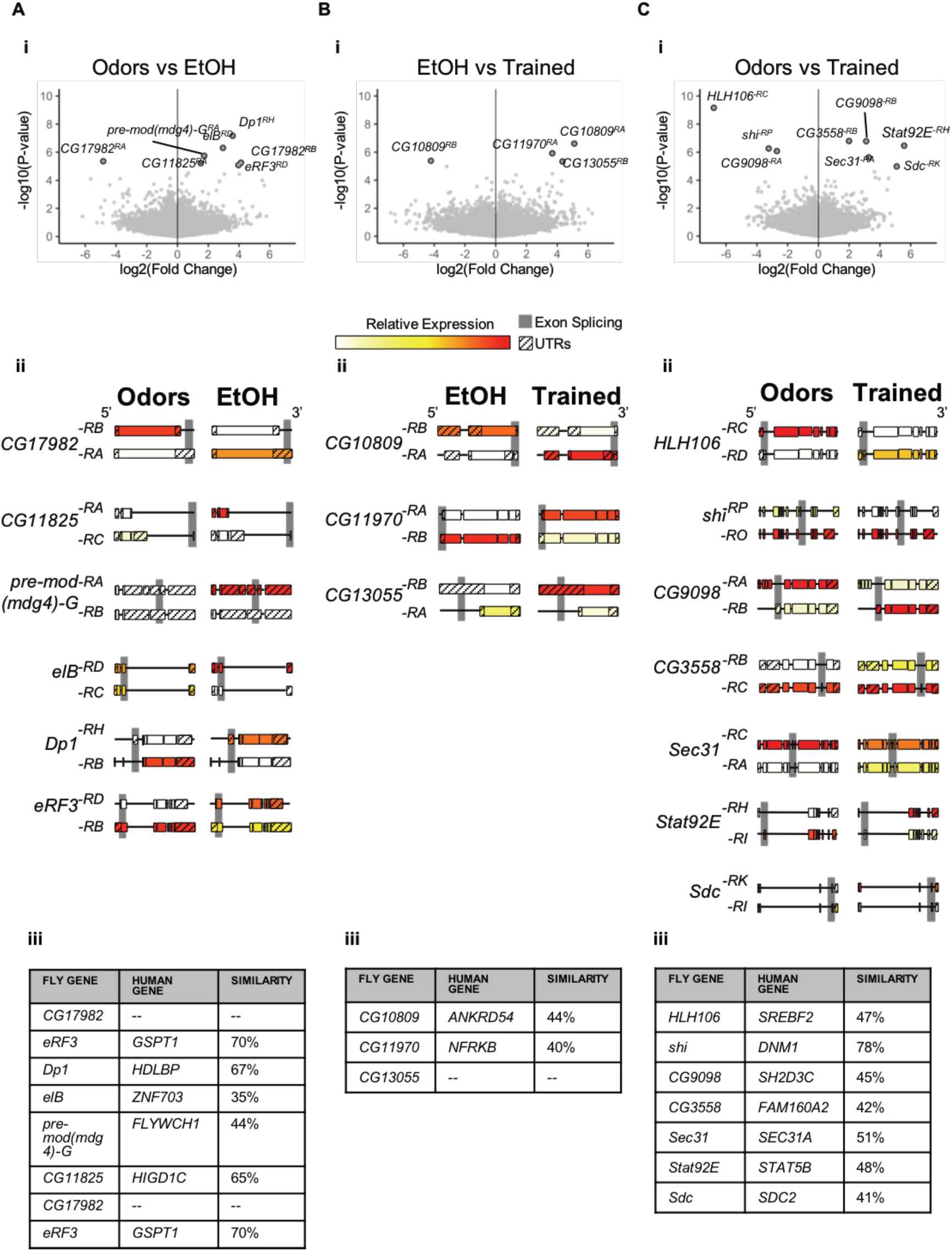
Differential transcripts across particular pairwise treatment comparisons. **(i)** Volcano plots of **(A)** Odors vs EtOH, **(B)** EtOH vs Trained and **(C)** Odors vs Trained comparisons (dark outline, FDR < 0.05). **(ii)** Significantly altered transcripts are depicted alongside another highly expressed isoform of the same gene, with gray shaded regions denoting splicing differences between isoforms. **(iii)** Transcripts are listed in a table along with corresponding high DIOPT scoring human genes.

We found four transcripts (three genes, all of unknown function) that were differentially expressed in response to Trained treatment as compared to Ethanol alone (Figure 5B). One transcript was downregulated – *CG10809*^*-RB*^ – and three were upregulated – *CG11970*^*-RA*^, *CG13055*^*-RB*^, *CG10809*^*-RA*^.

We found eight transcripts (seven genes) that were differentially expressed in response to Trained treatment as compared to Odors only controls (Figure 5C, Supplemental Figure 3). Three of these transcripts were downregulated – the Sterol Regulatory Element Binding Protein SREBP (*HLH106*^*-RC*^), the *Drosophila* homolog of dynamin *shibire* (*shi*^*-RP*^), and a predicted Ras guanine-nucleotide exchange factor *CG9098*^*-RA*^. Four transcripts were upregulated – a retinoic acid-like protein *CG3558*^*-RB*^, the same predicted Ras guanine-nucleotide exchange factor *CG9098*^*-RB*^, a Coat Protein Complex II factor *Secretory 31* (*Sec31*^*-RA*^), a transmembrane heparin sulfate proteoglycan *Syndecan* (*Sdc*^*-RK*^), and a transcription factor that functions in the JAK/STAT pathway *Signal-transducer and activator of transcription protein at 92E* (*Stat92E*^*-RH*^). The predicted splice junctions associated with these transcripts suggest a complex pattern of expression (Supplemental Figure 6). Using the DRSC Integrative Ortholog Prediction Tool (DIOPT) (Hu *et al*. 2011) we identified, when possible, the human genes corresponding to the implicated fly genes (Figure 5Aiii,Biii,Ciii).

### Molecular interactions with candidate ethanol cue memory genes

To better visualize and probe deeper into the known functions of differentially expressed transcripts associated with formation of ethanol cue memories, we used the open source Cytoscape platform (Shannon *et al*. 2003) and MIST database (Hu *et al*. 2018) to identify proteins that associate or interact with Odor vs Trained genes of interest (Figure 6). By overlaying our dataset information like expression level and fold change, onto this network, we provide a foundation for prospective hypothesis-driven bioinformatic inquiries and experiments. Notably, Stat92E had the largest known network, including a number of proteins associated with nucleosome remodeling (HDAC1, brm, mor), transcriptional regulation (Taf1, Ada2b, kay, Jra, CG13510), and splicing (Cdc5, CG7564, tsu,). Altogether the novel targets, and their interactors, offer experimental avenues to explore in future *Drosophila* and mammalian studies.

**Figure 6.**
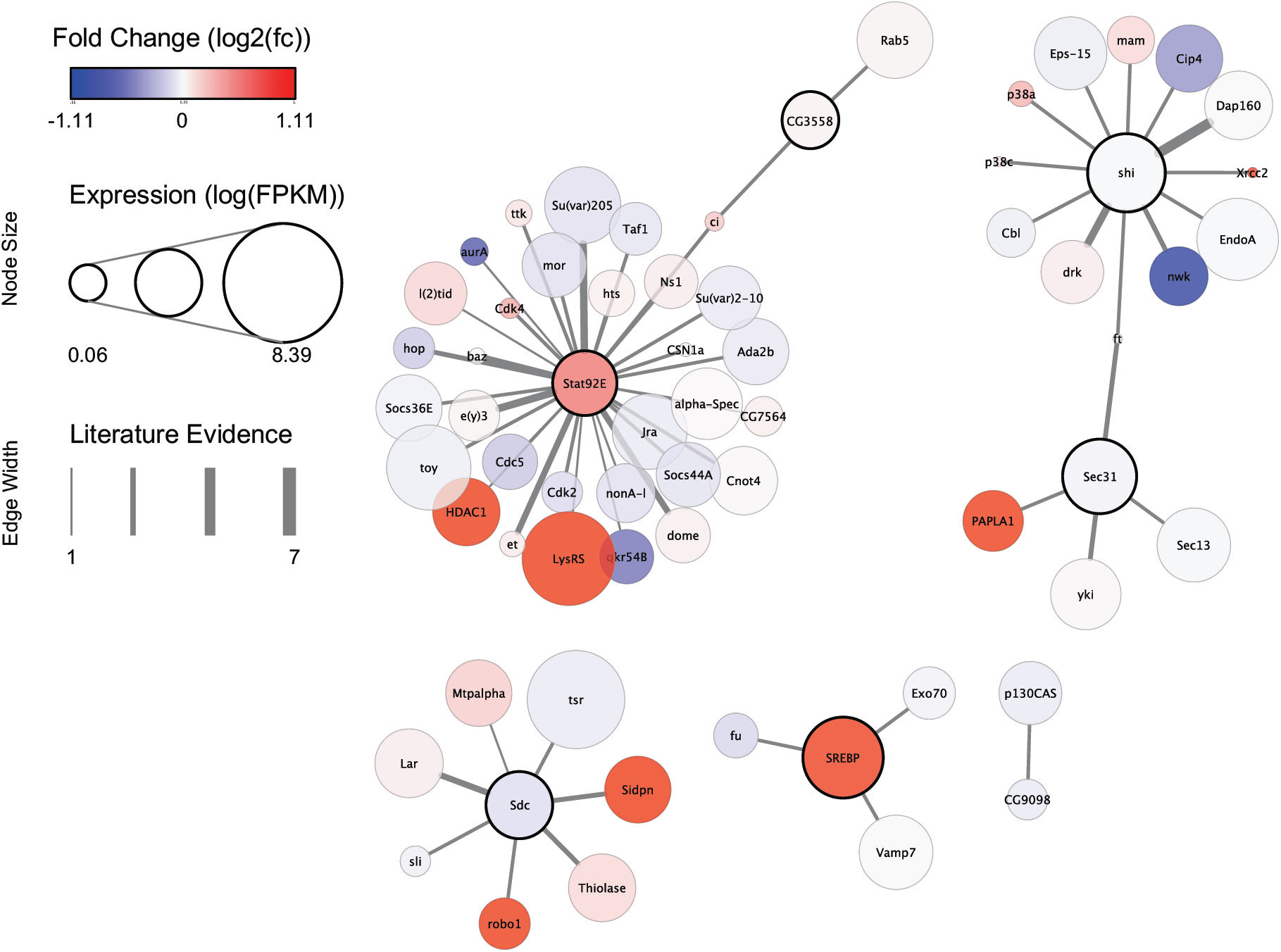
Protein interaction network of proteins associated with Odor vs Trained significant transcripts. Each circle node represents a protein, with dark outlines denoting proteins associated with significant Odor vs Trained transcripts. Attributes of the nodes includes fold change as color, expression level as size and edge thickness between nodes represents the extent of protein-protein interaction sevidence from the MIST database.

## Discussion

The nuclear and cell-type specific nature of our data provide a unique analysis of genes actively transcribed in the nucleus of memory-encoding neurons 24 hours after experience with alcohol or memory formation. Selection of poly-adenylated transcripts further allowed us to investigate changes to processed transcripts, rather than nascent RNA.

### Known genes implicated in alcohol experience and memory formation

Two of the genes that express alternative transcript isoforms in our ethanol cue memory data have been functionally implicated in similar behaviors. *shi*, the fly homolog of Dynamin (van der Bliek and Meyerowitz 1991) plays a role in endocytosis critical for synaptic transmission in MB neurons (McGuire *et al*. 2001; Kasuya *et al*. 2009) and in ethanol tolerance (Krishnan *et al*. 2012). We’ve also previously shown that expressing a dominant negative form of *shi* in MB neurons disrupts ethanol cue memory (Kaun et al, 2011). Stat92E, a transcription factor in the JAK/STAT signaling pathway, plays a role in long-term memory in *Drosophila* (Copf et al. 2011). We’ve recently shown that decreasing Stat92E in MB neurons similarly reduces ethanol cue memory (Petruccelli *et al*. 2018). To our knowledge, very few gene targets isolated from this study have been implicated in long-term memory in *Drosophila* (Supplemental Figure 7). However, a recent study showing the effects of acute response to alcohol shows some overlap in transcriptomic targets (Supplemental Figure 7) (Signor and Nuzhdin 2018).

### Context specific transcript expression

An intriguing pattern that emerged from our data is that administration of repeated mild doses of alcohol, odors alone, or odors paired with ethanol all produced significantly altered transcript isoforms. Remarkably, the particular isoforms identified were specific to the behavioral paradigm the animals experienced. Even altered transcripts associated with the presentation of alcohol alone were very different from those when alcohol was presented concomitantly with an odor. This suggests that transcript isoform expression is specific to the type of memory formed.

### Experience-dependent alternative splicing in MB neurons

Alternative transcript isoforms can be expressed through alternative transcription start sites or alternative splicing. Intriguingly, there are known protein-protein interactions between Stat92E and spliceosome complex associated proteins, including Cdc5, CG7564, and tsu (Guruharsha *et al*. 2011) (Figure 6). This suggests that splicing may readily occur in MB neurons as a result of experience. A caveat to our data, however, is that because we did not *a priori* expect to see splicing differences, our sequencing data was not performed with long-read sequencing techniques. Nor were statistical considerations, such as k-means clustering of transcripts or weighting exon-exon junction reads, used to compare splicing across genes.

The most direct cause of abnormal splicing is due to a mutation in the core splicing consensus sequences (Cieply and Carstens 2015). Our data, however, are derived from animals with the same genetic backgrounds and rearing, and demonstrate very few altered transcripts are detected 24 hours after experience. This suggests that mutagenesis is not the cause of the potential splicing differences we observed. Furthermore, although our data showed no direct evidence that spliceosomal machinery was differentially expressed with treatment, this does not rule out the possibility that spliceosome-associated proteins were acutely affected during treatment exposure. Our data may represent the aftermath of these, or post-translational response, effects.

### Splicing associated with memory formation

Splicing of particular targets in response to memory formation is not without precedent. For example, the splicing of the *Orb2A* transcript is required for both associative appetitive and courtship suppression memory in *Drosophila* (Gill *et al*. 2017). There is also evidence that this phenomenon may be conserved across species, since contextual fear conditioning induces differential splicing in the hippocampus of mice (Poplawski *et al*. 2016). It is, therefore, conceivable that whereas splicing may occur during all forms of synaptic plasticity, the isoform targets may be specific to the type of memory formed. The broad implication of this prediction is that the molecular engram of memory within relevant cells uses transcriptional diversity to provide the cellular plasticity associated with the type of memory being formed. This cellular encoding is bolstered by the diversify of epigenetic changes caused by neuronal activity associated with memory formation. For example, activity-induced histone modifications caused a late-onset shift in Neurexin-1 splicing reduced the stability of memories (Ding *et al*. 2017). Our data suggest that alcohol can affect transcriptional events, and thus shape how context is encoded during formation of memories.

### Alcohol regulates splicing in cue-encoding neurons

Several recent studies have shown that alternative splicing is correlated to chronic alcohol consumption. Chronic self-administration of alcohol in cynomolgus macaques (*Macaca fascicularis*) is associated with alternative splicing of AMPA subunits in the prefrontal cortex (Acosta *et al*. 2011). Broader transcriptomic analysis demonstrated over-representation of genes associated with splicing across brain regions in chronic ethanol self-administrating rhesus macaques (*Macaca mulatta*) (Iancu *et al*. 2018). Similar alcohol-induced effects on splicing extend to humans (Farris and Mayfield 2014). Post-mortem analysis of the brains of AUD patients showed novel splicing in *GABAB1* that decreased expression of the GABA binding site (Lee *et al*. 2014). Similarly, alcohol-induced splicing events also occurred in the developing human cortex *in utero*, potentially resulting in devastating neurodevelopmental consequences (Kawasawa *et al*. 2017).

Our data suggest that repeated alcohol exposures have lasting consequences on the transcript isoforms expressed in *Drosophila*. Importantly, altered isoform transcription occurred in a diversity of genes, including transcription factors like Stat92E, which could have long-lasting and broad effects on transcription within memory circuits (Copf *et al*. 2011). As alcohol alone did not induce expression of alternative transcripts of Stat92E, this suggests that splicing of this gene might occur due to memory formation. Since our analysis was restricted to neurons necessary for memories associated with punishment or reward, it is possible that even small effects could have widespread consequences for subsequent memory formation and vulnerability to dependence on drugs of abuse. Uncovering whether this is a broader phenomenon will require identifying lasting transcriptional events that occur in the same cell types across a diversity of memory and drug exposure paradigms.

### Diverse splicing mechanisms across addictions

Alternative splicing has been recently implicated in cocaine addiction (Cates *et al*. 2018). Transcription factor E2F3a regulates cocaine-induced alternative splicing in the mouse nucleus accumbens, and E2F3b mediates cocaine responses in the prefrontal cortex (Cates *et al*. 2019). *Drosophila E2F1* was expressed in our MB nuclei dataset (Figure 2cii), although it was not alternatively spliced in response to alcohol or alcohol-cue training (See transcript count data in GEO). We speculate that cocaine utilizes the E2F transcription factor family more than alcohol does to drive drug-cue memory formation and retention. Future investigations will likely identify both conserved drug-specific and convergent molecular mechanisms that influence transcriptional activity in reward circuitry.

## End Materials

### Data Availability

Raw sequencing data and count data presented in this work will be made freely available on NCBI Gene Expression Omnibus via accession number GSE108525.

### Author Contributions and Notes

E.P. and K.R.K. designed the research and wrote the manuscript. E.P. and N.L. performed INTACT nuclear extraction. N.L. developed the pipeline to align RNA-seq reads. E.P. performed statistical analyses and produced figures in consultation with K.R.K. Revision of the manuscript was performed by E.P., N.L., and K.R.K.

The authors declare no conflict of interest.

Supporting information is addended following references of this article.

## Acknowledgments

We thank Lee Henry (CSHL) for advice on INTACT, and members of the Kaun Lab, especially Reza Azanchi, Michael Feyder, and Yanabah Jaques for assistance with *Drosophila* husbandry and lab tasks. We thank Dr. Faith Liebl (SIUE), Dr. Kate O’Connor-Giles (Brown University), and members of the Kaun lab for helpful feedback on the manuscript. This work was supported by the Richard and Susan Smith Family Foundation, Newton, MA, the Carney Institute for Brain Science Center of Biomedical Research Excellence “Center for Nervous System Function” (NIGMS P20GM103645), and the National Institute on Alcohol Abuse and Alcoholism (R01AA024434).

## Supplemental Information

**Supplemental Figure 1.**
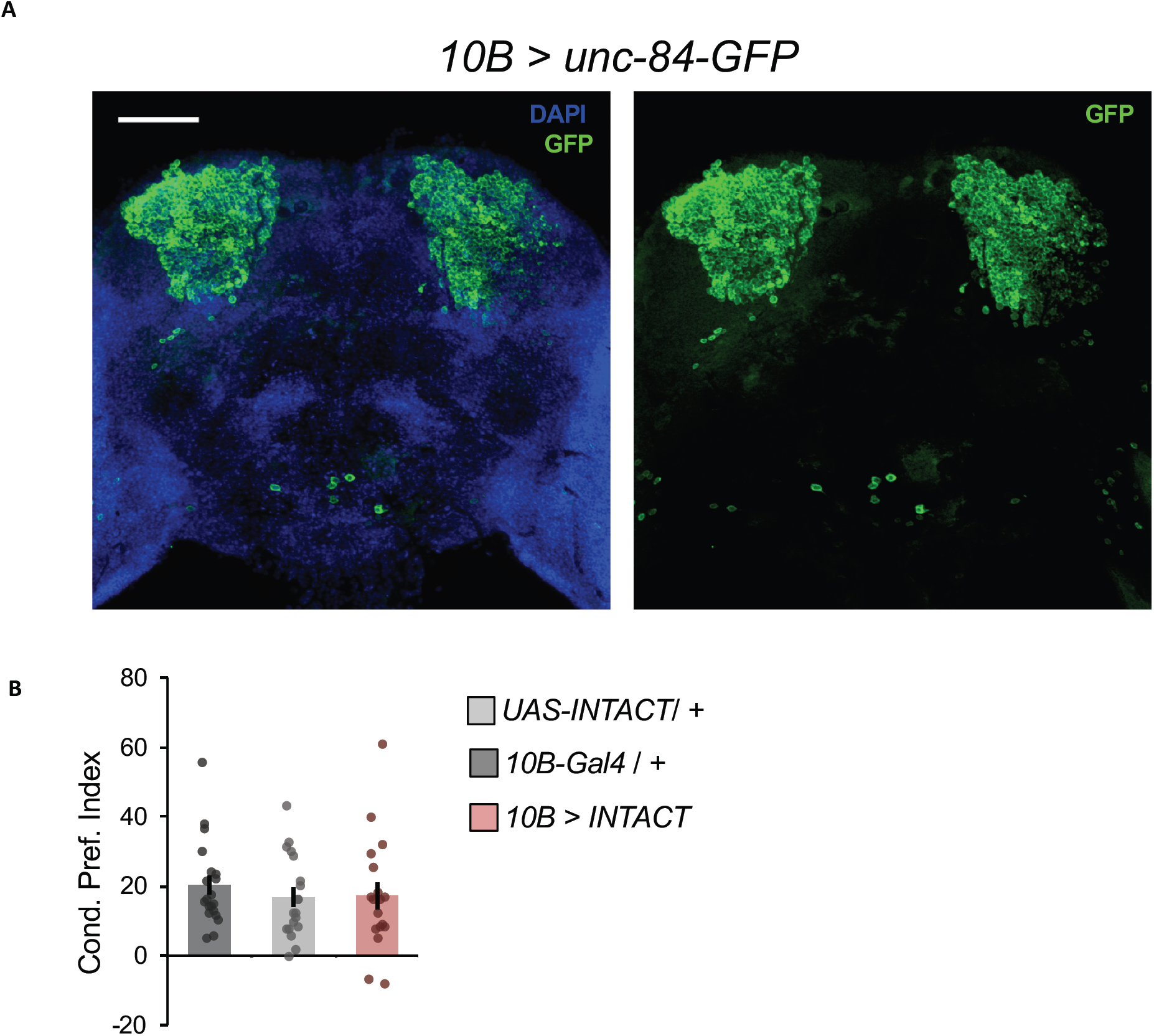
Restricted expression of INTACT transgene to all MB nuclei. **A)** A representative immunohistochemical staining of a 3-5-day-old adult male brain (DAPI staining DNA, Blue; anti-GFP staining GFP, Green). Confocal max stack image taken at 20x resolution (Scale bar 50 um). **B)** Expression of the INTACT transgene (unc^84^-GFP) in MB neurons did not interfere with alcohol associative preference; Gal4 control (n = 19), UAS control (n= 20), Gal4>UAS experimental line (n= 18).

**Supplemental Figure 2.**
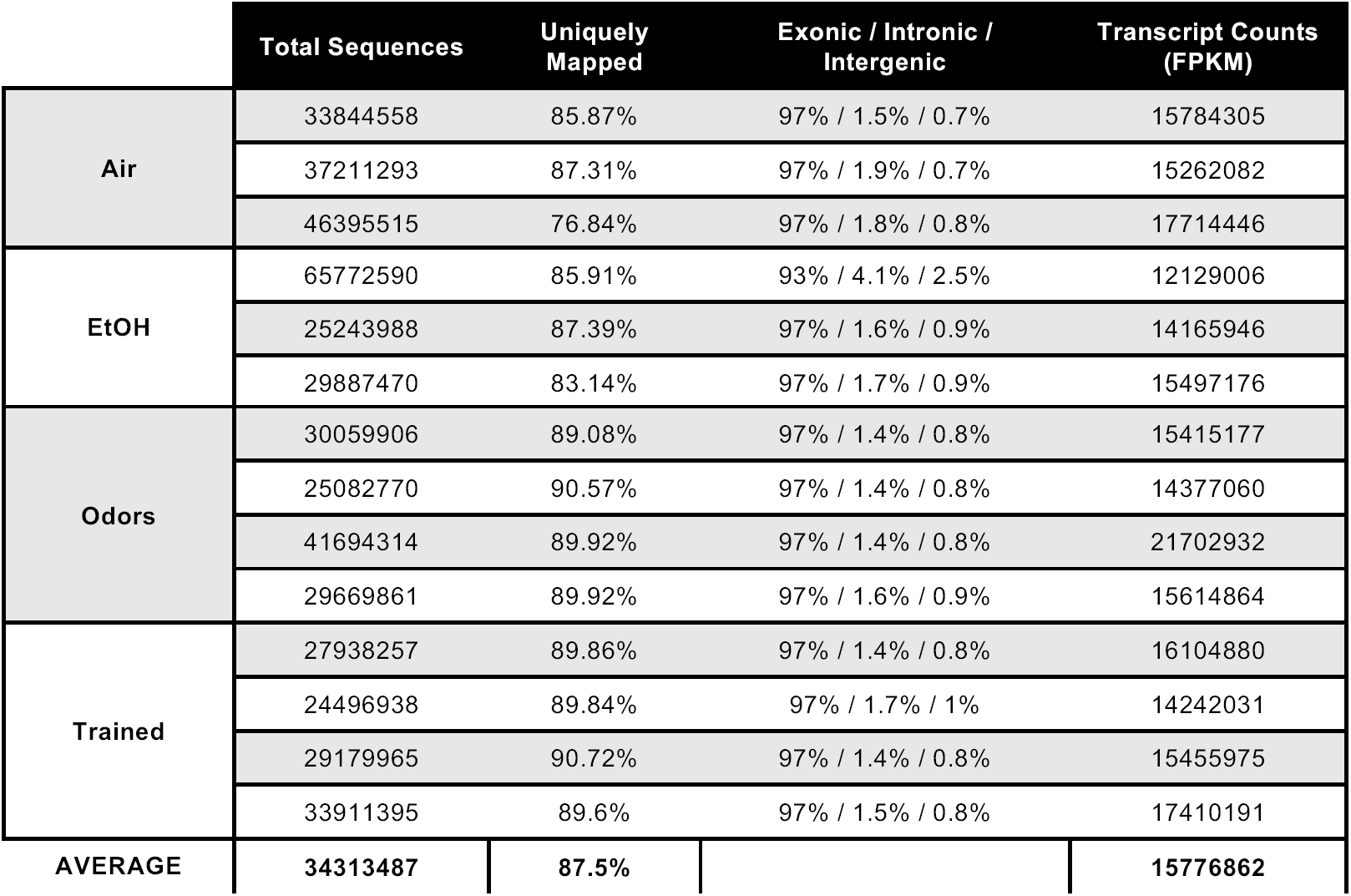
QualiMap (BAM QC) analysis of the RNA-seq libraries. The total sequences, uniquely mapped reads %, genomic biotype, and normalized transcript counts (FPKM) are displayed for all 14 libraries.

**Supplemental Figure 3.**
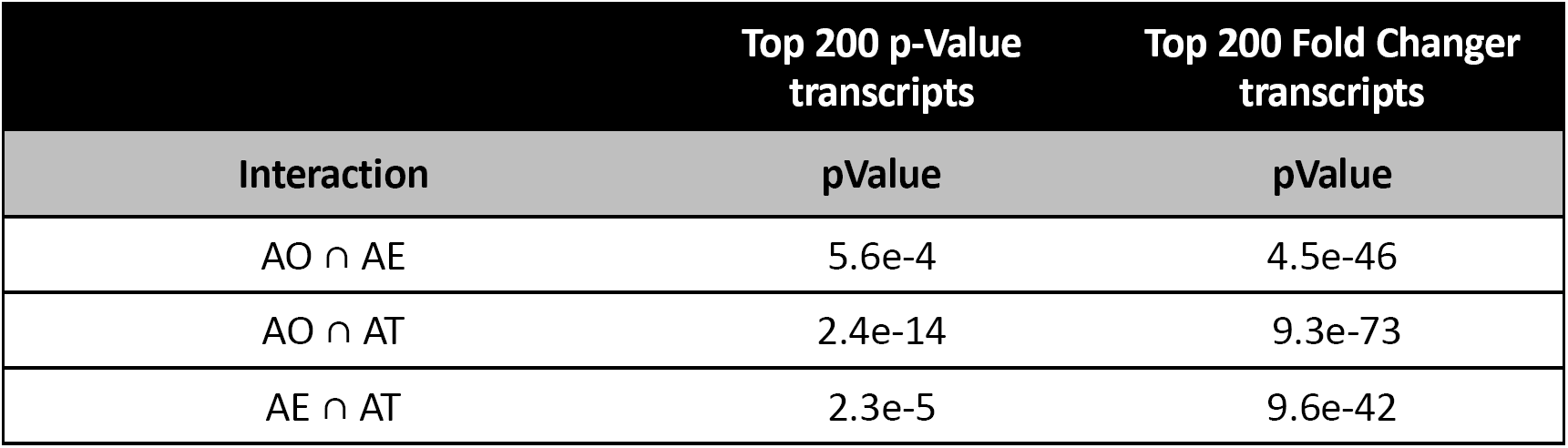
Hypergeometric distribution test p-values. The phyper function in R was used to determine hypergeometric p-values for data in text Figure 4B,E. The ‘universe’ test number was 34,741 transcripts from the current Dm6 transcriptome. Values are as follows: phyper(q, m, n, k, lower.tail=FALSE) … eg. phyper(overlap#, 200, (34741-200), 400, lower.tail=FALSE).

**Supplemental Figure 4.**
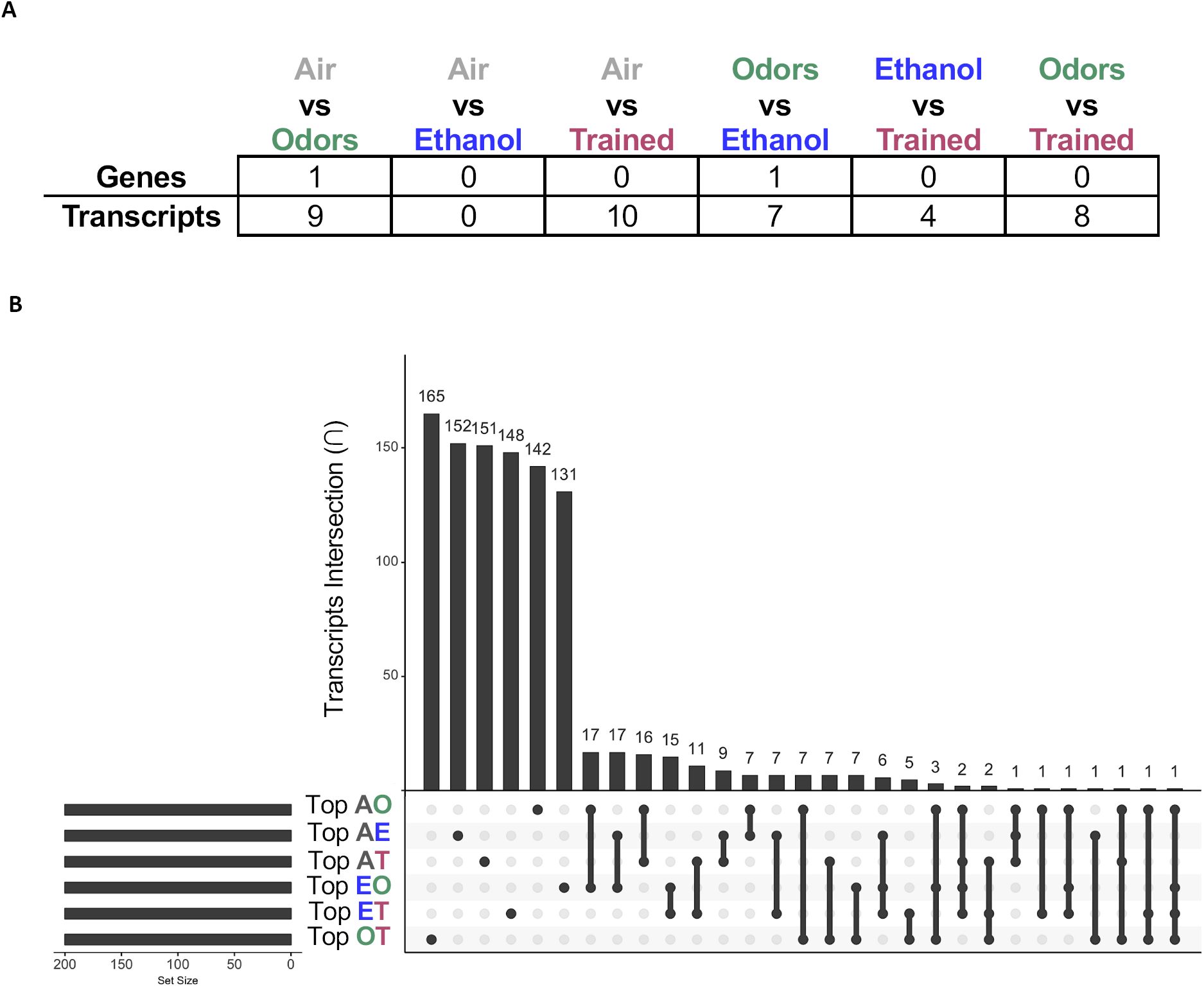
Intersectional analysis of top differentially expressed transcripts across treatments, as compared to Air Controls. **A)** A table representing the number of genes or transcript reaching FDR < 0.05 between all pairwise comparisons: Air vs Odors, Air vs EtOH, Air vs Trained, Odors vs EtOH, EtOH vs Trained, and Odors vs Trained. Colors depict treatment condition. **B)** An upgraded Venn Diagram plot generated by an R package called ‘UpSetR’ demonstrating the intersection between top 200 p-value transcripts (abbreviated by first letter in treatment).

**Supplemental Figure 5.**
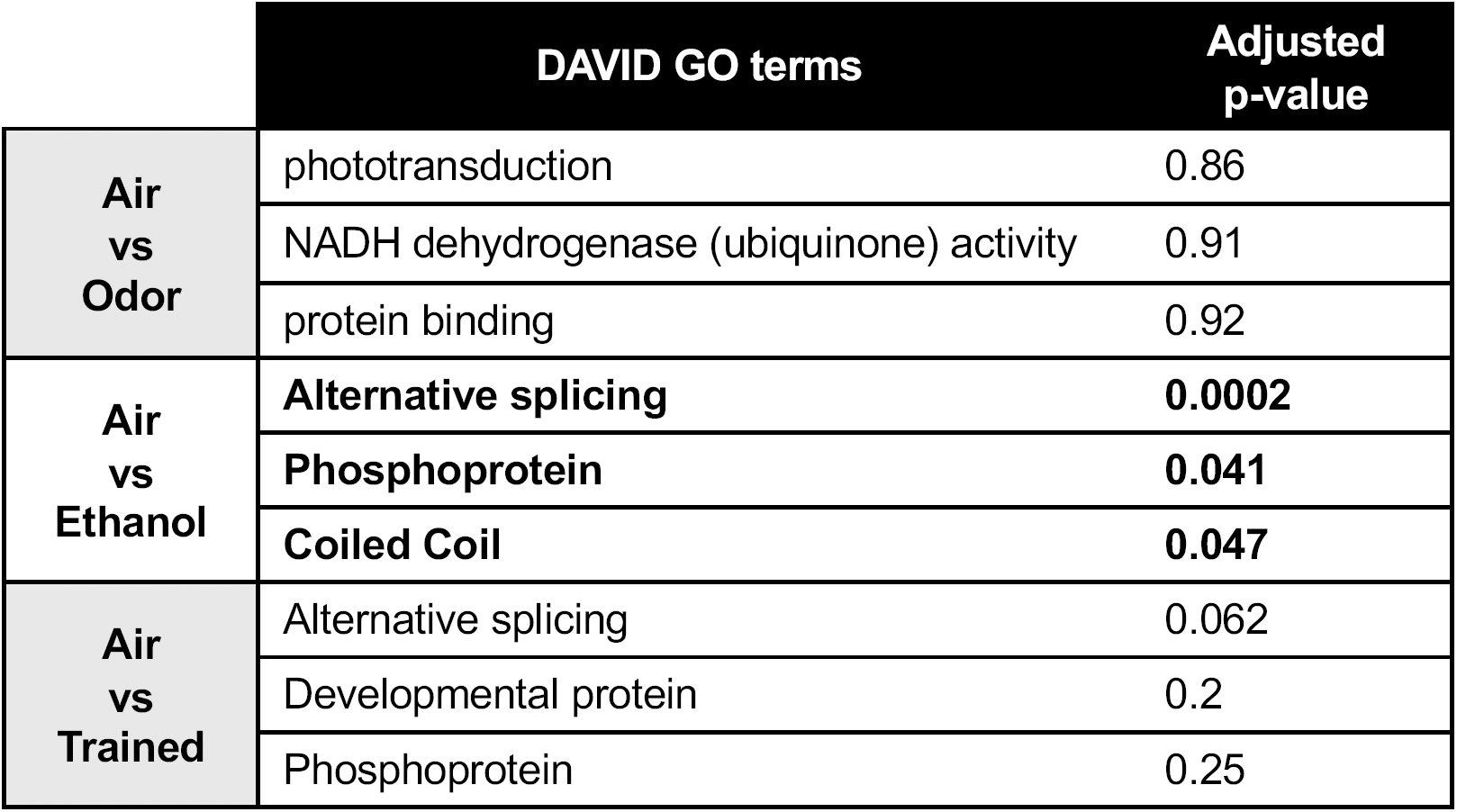
DAVID analysis of gene ontology (GO) of top 200 p-value genes. Enrichment of current GO terms within the top 200 p-value genes found between Air vs Odor, Air vs Ethanol, and Air vs Trained pairwise comparisons (Bolded adjusted p-value < 0.05).

**Supplemental Figure 6.**
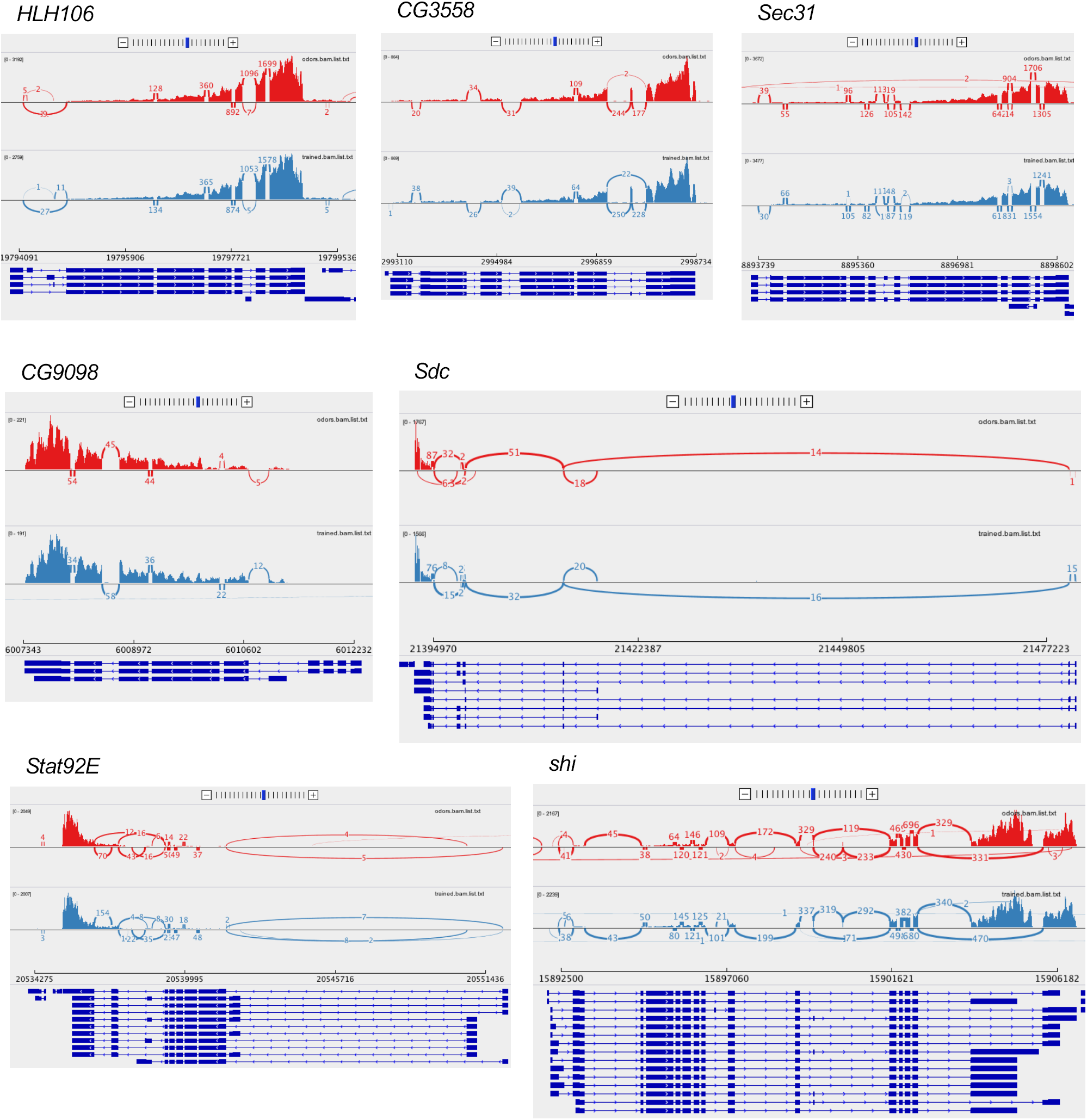
Sashimi plots from IGV (Broad Institute). Sashimi plots help to visualize splice junctions across merged ‘Odors’ and ‘Trained’ samples for differentially expressed transcripts.

**Supplemental Figure 7.**
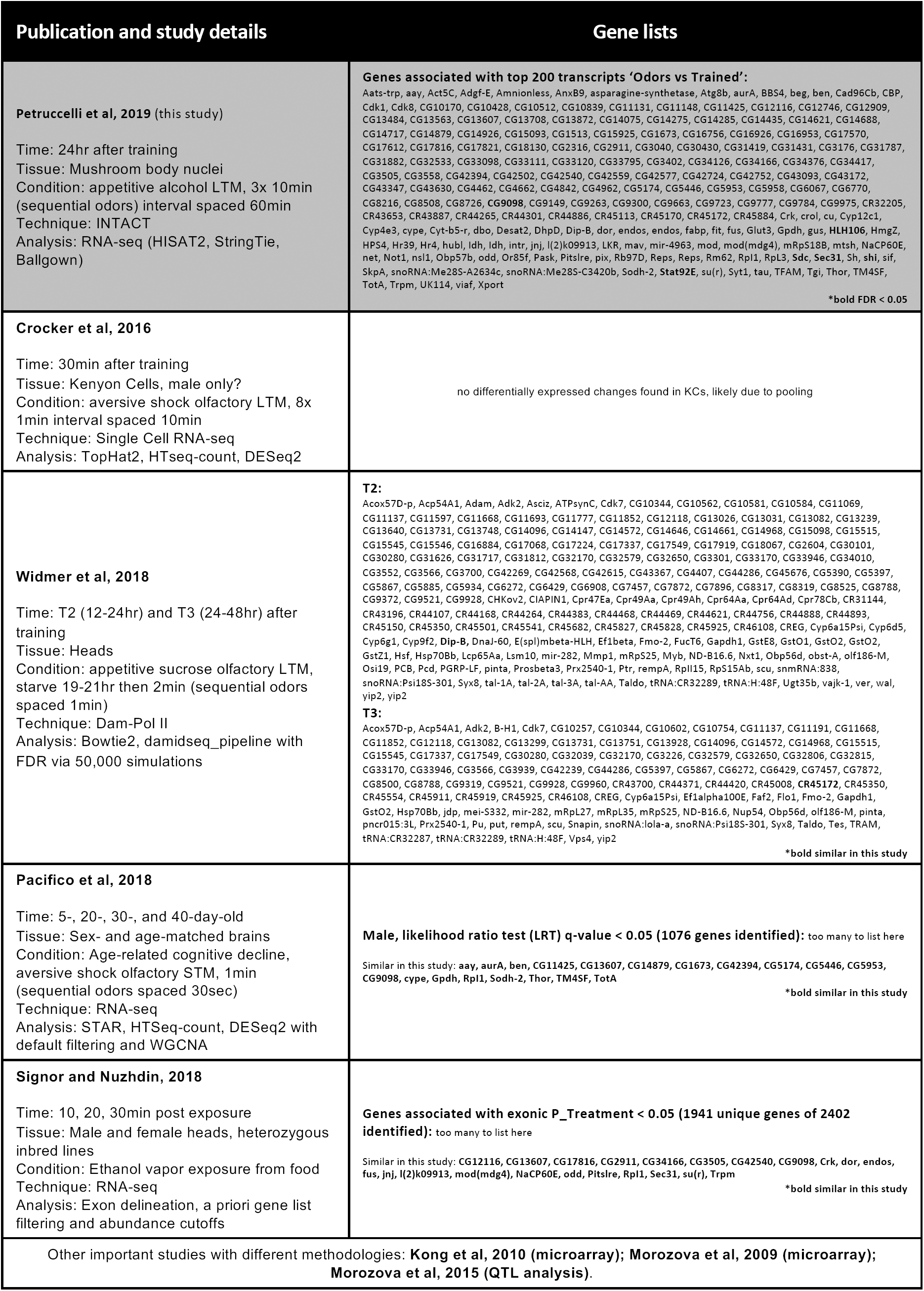
Comparison of our results to previous RNA-Seq studies.

